# Chromatin accessibility landscape of articular knee cartilage reveals aberrant enhancer regulation in osteoarthritis

**DOI:** 10.1101/274043

**Authors:** Ye Liu, Jen-Chien Chang, Chung-Chau Hon, Naoshi Fukui, Nobuho Tanaka, Zhenya Zhang, Ming Ta Michael Lee, Aki Minoda

## Abstract

**Background:** Osteoarthritis (OA) is a common joint disorder with increasing impact in an aging society; however, there is no cure or effective treatments so far due to lack of sufficient understanding of its pathogenesis. While genome-wide association studies (GWAS) and DNA methylation profiling identified many non-coding loci associated to OA, the interpretation of them remains challenging.

**Methods:** Here, we employed Assay for Transposase-Accessible Chromatin with high throughput sequencing (ATAC-seq) to map the accessible chromatin landscape in articular knee cartilage of OA patients and to identify the chromatin signatures relevant to OA.

**Results:** We identified 109,215 accessible chromatin regions in cartilage and 71% of these regions were annotated as enhancers. We found these accessible chromatin regions are enriched for OA GWAS single nucleotide polymorphisms (SNPs) and OA differentially methylated loci, implying their relevance to OA. By linking these enhancers to their potential target genes, we have identified a list of candidate enhancers that may be relevant to OA. Through integration of ATAC-seq data with RNA-seq data, we identified genes that are altered both at epigenomic and transcriptomic levels. These genes are enriched in pathways regulating ossification and mesenchymal stem cell (MSC) differentiation. Consistently, the differentially accessible regions in OA are enriched for mesenchymal stem cell-specific enhancers and motifs of transcription factor families involved in osteoblast differentiation (e.g. bZIP and ETS).

**Conclusions:** This study marks the first investigation of accessible chromatin landscape on clinically relevant hard tissues and demonstrates how accessible chromatin profiling can provide comprehensive epigenetic information of a disease. Our analyses provide supportive evidence towards the model of endochondral ossification-like cartilage-to-bone conversion in OA knee cartilage, which is consistent with the OA characteristic of thicker subchondral bone. The identified OA-relevant genes and their enhancers may have a translational potential for diagnosis or drug targets.

## Background

Osteoarthritis (OA) is a degenerative joint disease [1,2] that is one of the most common causes of chronic disability in the world [3,4], of which the knee OA is the most common. Main features of OA include cartilage degradation, subchondral bone thickening, joint space narrowing and osteophytes formation [5], resulting in stiffness, swelling, and pain in the joint. Currently available treatments are either pain relief or joint function improvement by strengthening the supporting muscles. However, OA progression ultimately leads to costly total joint replacement surgery, making it a growing global health burden.

Although the causes of OA are not well understood, risk factors such as age, weight, gender, and genetic factors have been identified [4]. Several models for OA initiation, such as mechanical injury, inflammatory mediators from synovium, defects in metabolism and endochondral ossification, have been proposed to explain pathogenesis of this disease [6–12]. To date, GWAS have identified more than 20 loci to be associated with the risk of developing OA [13]. While next generation sequencing data is being generated to discover rare variants with larger effect size, the identified variants are often located in the non-coding regions of the genome [14], complicating the identification of the causal genes. Transcriptomic analyses of cartilage in diseased joints of OA patients (taken from the replacement surgeries) provided an opportunity to pinpoint transcriptionally dysregulated genes and pathways relevant to OA [15,16]. However, such studies have yet to fully reveal the underlying molecular mechanism of how the transcription of these genes are dysregulated.

Recently, epigenetic tools have been applied to gain further insight into the pathogenesis of OA. There have been several reports of DNA methylation status [17–19] in cartilage of diseased joints. However, relatively few DNA methylation alterations were found in OA degenerated tissues, despite many genes being differentially expressed. Consistent with these findings, it was also found that change of gene expression is rarely associated with DNA methylation alterations at promoters [16,19,20]. Many of the identified differentially methylated sites in fact fall in enhancer regions, which are non-coding regulatory elements, disruption of which may lead to dysregulated transcription, and many are cell type-specific [21,22]. Recent large-scale studies, such as FANTOM5 [23], Roadmap Epigenomics Project [24] and GTEx [25] have enabled the prediction of regulatory networks between enhancers and their potential target genes (e.g. JEME [26]), which could be applied in a clinical context to explore the roles of enhancers in disease pathogenesis.

Here, we set out to investigate alterations of enhancers associated with OA by applying ATAC-seq [27] on the knee joint cartilages from OA patients, using an optimized protocol for cartilage sample preparation. ATAC-seq maps the accessible chromatin regions, which are often regulatory regions such as promoters and enhancers that play roles in regulation of gene expression. By integrating our ATAC-seq data with the publicly available genetic, transcriptomic and epigenomic data, we identified dysregulated enhancers and their potential target genes. Our data highlights a number of OA risk loci and differentially methylated loci that potentially play roles in cartilage degradation during OA development.

## Methods

### Knee joint tissues collection

Human knee joint tibial plates were collected from patients who undertook joint replacement surgery due to severe primary OA at the National Hospital Organization Sagamihara Hospital (Kanagawa, Japan). Demographic information for patients ism listed in Additional file 1: Table S1. Diagnosis of OA was based on the criteria of the American College of Rheumatology [28], and all the knees were medially involved in the disease. Tissues were stored in 4°C cultured medium after removal. Informed consent was obtained from each patient enrolled in this study. This study was approved by the institutional review board of all the participating institutions (Sagamihara National Hospital and RIKEN).

### Processing of cartilage samples

Fresh cartilage was separated from the subchondral bone by a scalpel immediately after surgery and stored in 4°C Dulbecco’s Modified Eagle Medium until they were processed. 100 mg cartilage from both outer region of lateral tibial plateau (oLT) and inner region of medial tibial plateau (iMT) (Fig. 1a and Additional file 2: Figure S1a) for each patient was digested with 0.2% type II collagenase (Sigma-Aldrich) in DMEM at 37°C with rotation for 12 hours to fully remove the collagen matrix debris, and stopped by washing with chilled PBS. Immediately after digestion, chondrocytes were counted by a cell counting chamber Incyto C-Chip (VWR) and collected by centrifugation at 500g for 5 minutes at 4°C.

**Fig. 1.**
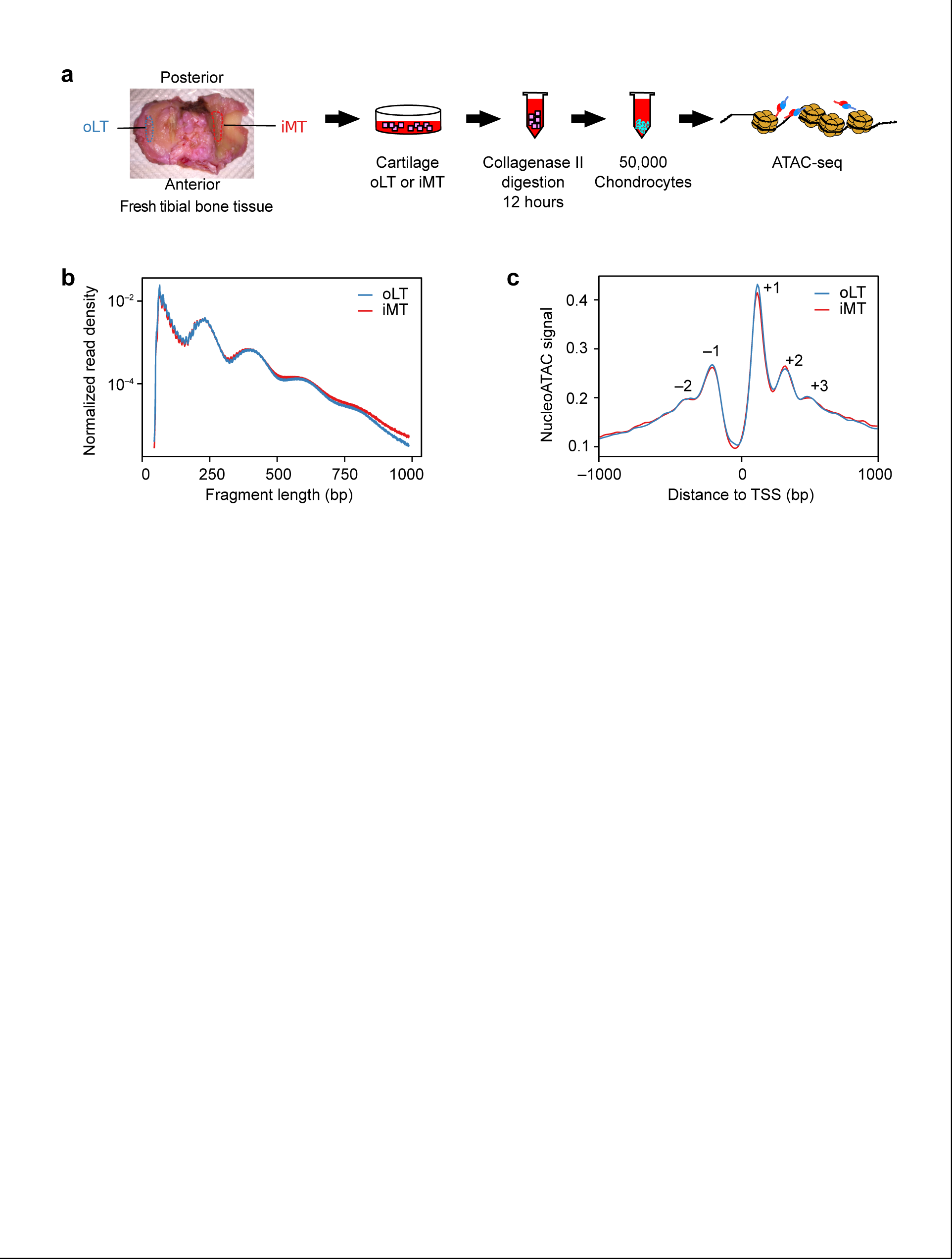
ATAC-seq of OA cartilage tissues. (a) Schematic diagram of the experimental flow. Chondrocyte from two regions of the articular cartilages were isolated for ATAC-seq. oLT (outer Libial Tibial): intact tissue. iMT (inner Medial Tibial): damaged tissue. (b) ATAC-seq sequencing fragment length distribution for pooled oLT (n=8) and iMT (n=8) libraries. The banding pattern, representing fragments from nucleosome free regions, mono-, di-, and tri-nucleosome, indicates a successful ATAC-seq experiment. (c) Normalized nucleoATAC signal aggregated over all genes shows distinct nucleosome positioning around transcription start sites (TSS). Positive and negative numbers indicate the TSS upstream and downstream nucleosomes.

### ATAC-seq of the cartilage

Approximately 50,000 cells were taken after the cartilage processing and used for ATAC-seq library preparation (Fig. 1a) according to the original protocol [27,29]. Briefly, transposase reaction was carried out as previously described [27] followed by 9 to 12 cycles of PCR amplification. Amplified DNA fragments were purified with QIAGEN MinElute PCR Purification Kit and size selected twice with Agencourt AMPure XP (1:1.5 and 1:0.5 sample to beads; Beckman Coulter). Libraries were quantified by KAPA Library Quantification Kit for Illumina Sequencing Platforms (KAPA Biosystems), and size distribution was inspected by Bioanalyzer (Agilent High Sensitivity DNA chip, Agilent Technologies). Library quality was assessed before sequencing by qPCR enrichment of a housekeeping gene promoter (*GAPDH*) over a gene desert region [29]. ATAC-seq libraries were sequenced on an Illumina HiSeq 2500 (50bp, paired-end) by GeNAS (Genome Network Analysis Support Facility, RIKEN, Yokohama, Japan).

### ATAC-seq data processing, peak calling and quality assessment

An ATAC-seq data processing pipeline for read mapping, peak calling, signal track generation, and quality control was implemented [30]. Briefly, fastq files for all patients were grouped by tissue compartment (oLT or iMT) and input into the pipeline separately, with parameters *–true_rep –no_idr*. Reads were mapped to the hg38 reference genome. Peaks were called by macs2 with default parameters in the pipeline. Basic sequencing information and library quality metrics are listed in Additional file 1: Table S2. NucleoATAC was applied to infer genome-wide nucleosome positions and occupancy from the ATAC-seq data [31]. Briefly, NucleoATAC were run for oLT and niMT separately with default parameters, using bam files merging from all patients as input, and the outputted nucleoatac_signal.bedgraph were used for aggregated plot around TSS.

Previously annotated DNase I Hypersensitive Sites (DHS) were used to assess the quality of our libraries. Briefly, a set of DHS defined in Roadmap Epigenomics Project [32] were downloaded [33] and lifted over from hg19 to hg38 using UCSC liftOver tool [34] (refer as Roadmap DHS thereafter). For each library, we calculated the ratio of reads that fall within Roadmap DHS versus randomly chosen genomic regions of the same total size (DHS enrichment score). It is noted that the DHS enrichment score is in good correlation with enrichment qPCR for *GAPDH* (Additional file 2: Figure S1b).

### Defining a set of unified accessible chromatin regions

Peaks from all 16 libraries were pooled and those that are within 300 bp were merged, resulting in 615,454 merged raw peaks. Reads fall in raw peak regions were counted for individual samples using bedtools v2.27.0 [35]. Read counts were normalized as count-per-million (CPM) based on relative log expression normalization implementedm in edgeR v3.18.1 [36,37]. In the end, we defined a set robust peaks (n=109,215) with ≥2 CPM in at least 6 oLT or 6 iMT samples for all downstream analyses.

### Peak annotation and differentially accessible peak identification

Peaks were annotated as promoters or enhancers based on their intersection with promoter or enhancer as defined in Roadmap DHS [24]. Noted that a peak was preferentially annotated as a promoter if it intersects both a promoter and an enhancer DHS. Differentially accessible peaks between damaged (i.e. iMT) and intact (i.e. oLT) tissues were identified using edgeR [36,37]. Briefly, DHS enrichment score of each sample were incorporated as continuous covariates in the design matrix of the generalized linear model implemented in edgeR [36,37]. The Benjamini-Hochberg adjusted p-values (i.e. false discovery rate, FDR) was taken to measure the extent of differential accessibility as used in Fig. 3b. A cutoff of FDR ≤ 0.05 was used to define differentially accessible peaks.

**Fig. 2.**
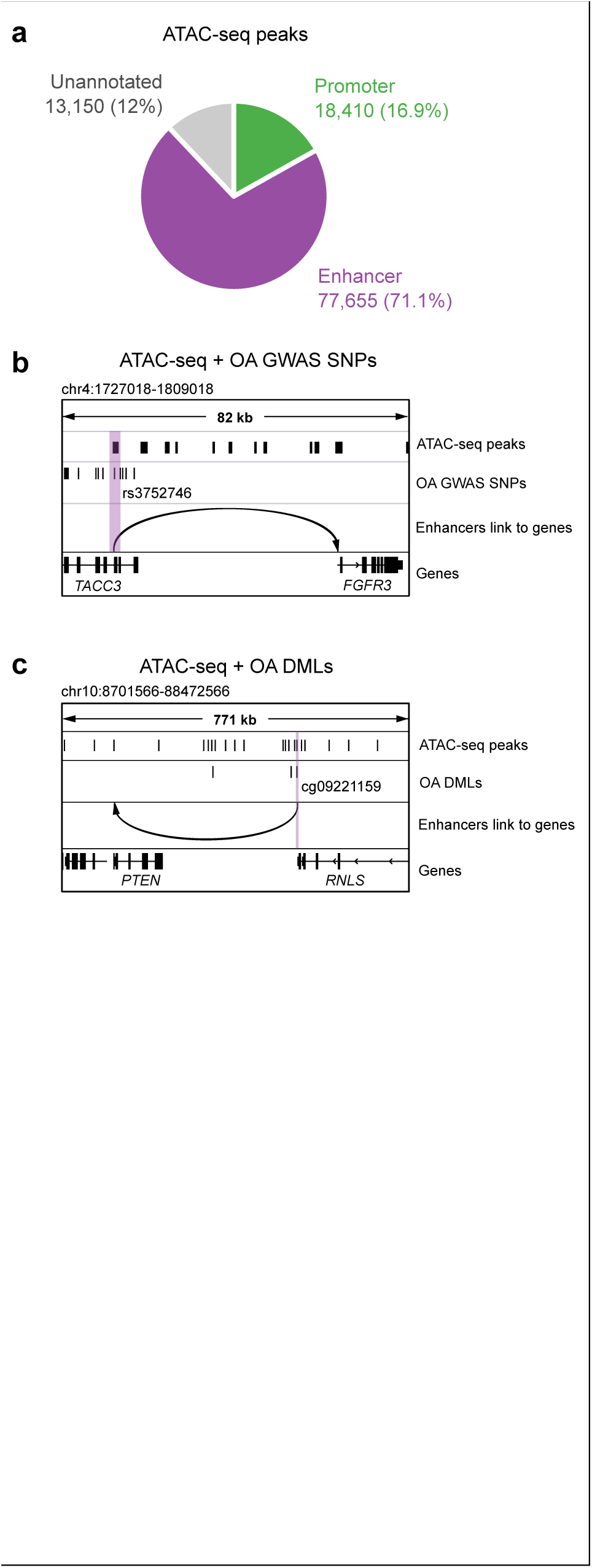
Identification of accessible chromatin regions with OA susceptible GWAS SNPs and differentially methylated loci. (a) Annotation of identified accessible chromatin regions. (b-c) An example of an OA GWAS SNP rs3752746 (b) and differentially methylated locus cg09221159 (c) overlapping with an open enhancer in cartilages from OA patients. The arrow points to the predicted target genes.

**Fig. 3.**
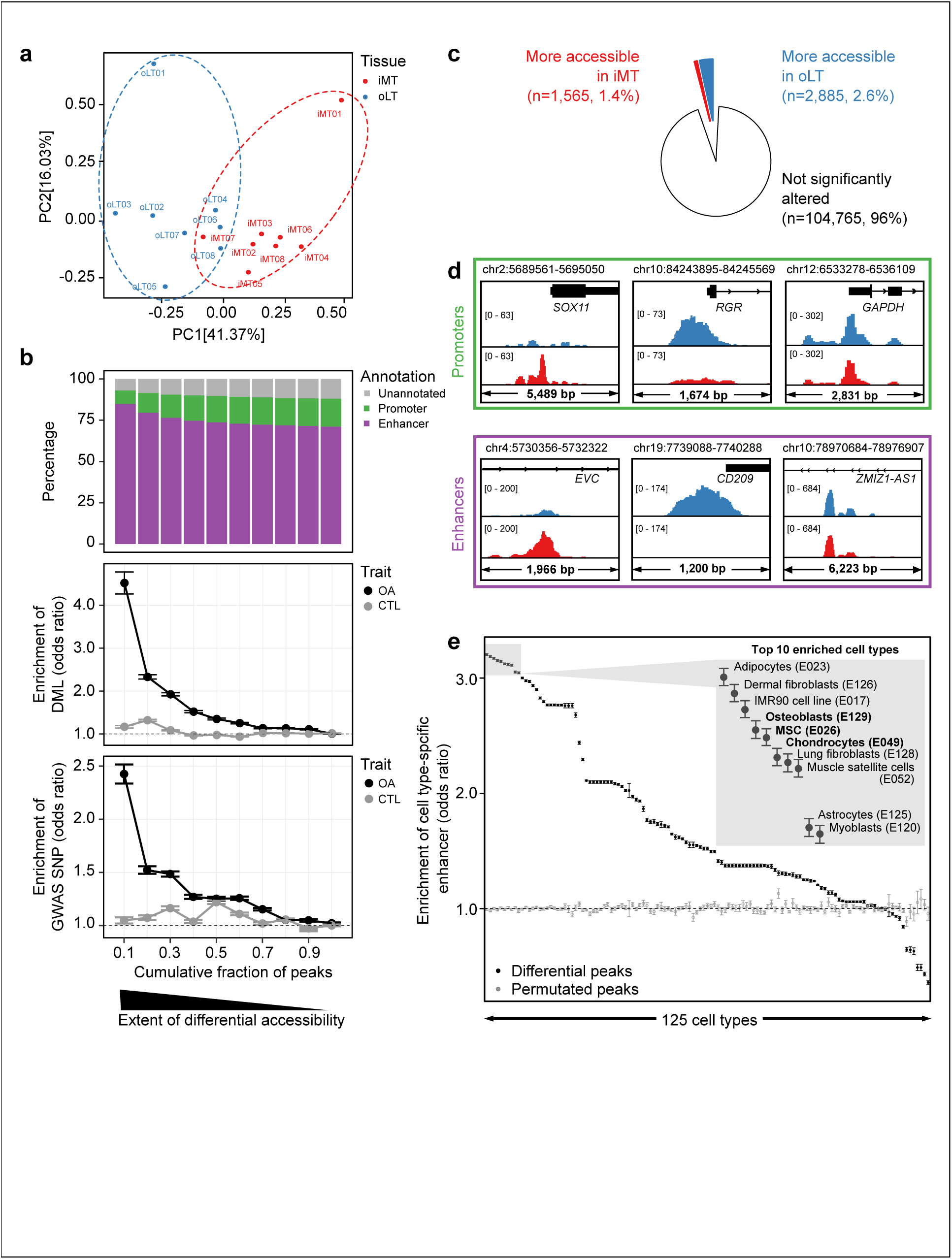
Bone-and chondrocyte-specific enhancers are dysregulated in OA cartilage. (a) Principal component analysis using ATAC-seq peaks shows general separation between oLT and iMT. (b) The extent of differential accessibility between iMT and oLT are correlated with enhancers composition (*top*), enrichment of OA differential methylated loci (DML, *middle*), and enrichment of OA GWAS SNPs (*bottom*). CTL: Control trait (Parkinson’s disease). Circles represent the mean and error bars represent the standard error for 100 times permutation. (c) Pie chart shows proportions of the ATAC-seq peaks that are more accessible in iMT (red) and oLT (blue). (d) Genome browser views of example loci (all patients pooled) showing differential accessible regions at promoters (*top*) and enhancers (*bottom*) with more accessible in iMT (*left*), more accessible in oLT (*middle*), and not significantly altered between oLT and iMT (*right*). (e) Enrichment of cell type-specific enhancers in differentially accessible peaks. Top 10 are listed (inset), which includes bone- and chondrocyte-related cell types (bold). Error bars represent 95% confidence interval.

### Processing of OA associated SNPs

GWAS lead SNPs of OA were obtained from GWASdb v2 (as of 16 Sep 2017) [38], NHGRI-EBI GWAS Catalog (as of 16 Sep 2017) [39], and a recent study [40]. SNPs in linkage disequilibrium with these lead SNPs (i.e. proxy SNPs with r^2^ > 0.5 andn distance limit of 500kb in any three population panels of the 1000 Genomes Project pilot 1 data) were obtained from SNAP database [41], resulting in 8,973 GWAS SNPs associated with OA. As a negative control, GWAS SNPs associated with Parkinson’s disease were obtained the same way from NHGRI-EBI GWAS Catalog. Coordinates for these SNPs in hg38 were obtained from dbSNP Build 150.

### Definition of cell type-specific enhancers

Bed files for enhancer clusters with coordinated activity in 127 epigenomes, as well as the density of clusters per cell type, were obtained from online database [33] (DNase I regions selected with p < 1 × 10^−2^, lifted over from hg19 to hg38 using UCSC liftOver tool). For each cell type, clusters with two-fold density than average (across all cell types) were defined as specific and pooled.

### Enrichment analysis for differentially accessible peaks

Fisher’s exact test were applied to assess the enrichment for the following features in the selected peaks: (1) GWAS SNPs associated with OA (n=8,973), (2) differentially methylated loci (DML) associated with OA (n=9,265) [19,42], and (3) cell type-specific enhancers. Briefly, for the enrichment of (1) GWAS SNPs and (2) DML, peaks were ranked by the extent of differential accessibility (as measured by FDR) and each cumulative fraction of peaks was compared to all peaks. GWAS SNPs and DML [43] associated with Parkinson’s disease was used as a negative control. For enrichment of (3) cell type-specific enhancers, differentially accessible peaks (FDR ≤ 0.05) were compared to all peaks, and same number of randomly selected peaks were used as a control. Permutation of peaks as control were done 25 times.

### Linking enhancer peaks to their potential target genes

A promoter peak is assigned to a gene if it intersects with the transcription start sites (TSS) of its transcript as defined in the FANTOM CAGE associated transcriptome [44]. We noted that a promoter peak might be associated with multiple genes. An enhancer peak is defined as linked to a target gene if it overlaps an expression quantitative trait loci (eQTL) of the corresponding gene (GTEx V7, in any tissues with p < 1 × 10^−5^) [45], or is supported by putative enhancer-promoter linkage predicted by JEME method [46] on FANTOM5 and Roadmap Epigenomics Project data [26].

### Integration with publicly available transcriptome data

We reanalyzed an RNA-seq dataset (ArrayExpress E-MTAB-4304) from an independent OA patient cohort, in which the RNA was extracted from cartilage tissues of both iMT and oLT for 8 patients [15]. Read counts on transcripts were estimated by Kallisto v0.43.1 [47] using default parameters on FANTOM CAGE associated transcriptome [44]. Estimated read counts of a gene, defined as the sum of the estimated read counts of its associated transcripts, were used as the input for differential gene expression analysis using edgeR v3.18.1. In total, 3,293 genes were defined as significantly differentially expressed between iMT and oLT (FDR ≤ 0.05). A gene is defined as “consistently dysregulated both at the epigenomic and transcriptomic levels” when it is upregulated (or downregulated) in RNA-seq with more (or less) accessible promoters or enhancers in ATAC-seq (n= 371).

### Gene ontology analysis

Enrichr [48] was used to identify gene sets enriched in genes implicated in OA. The selected gene lists were input to query enrichment for gene ontology (Biological Process 2017b) in the database. P-value ≤ 0.05 was considered significant.

### Transcription factor binding motif analysis

DNA motif analysis for differentially accessible regions was performed using HOMER v4.9 with default parameters [49]. The enriched *de novo* and known motifs as well as its matching transcription factor are searched and scanned in more- and less-accessible peaks separately (parameters: *-mask -size given*), with all robust peaks as background region. P-value < 1 × 10^−5^ was considered significant.

## Results

### Mapping chromatin accessibility of chondrocytes in OA knee cartilages

To investigate chromatin signatures in articular cartilage associated with OA, we performed ATAC-seq on the chondrocytes isolated from the knee joints of patients. We have previously shown that the oLT region (outer region of the lateral tibial plateau, representing the intact cartilage) is a good control for comparing the iMT region (inner region of medial tibial plateau, representing the damaged cartilage) as a model for OA disease progression [16], and the transcriptome and methylome of this model have been characterized by us and the others [19,20,50,51]. In this study, we performed ATAC-seq on the chondrocytes isolated from of the oLT and iMT regions of 8 patients (Additional file 2: Figure S1a), generating 16 ATAC-seq libraries (paired, from 8 patients) on clinically relevant OA cartilage tissues (Fig. 1a).

Overall, the libraries are of high quality, showing two-to four-fold enrichment in the Roadmap DHS (DHS enrichment score, Methods), with no substantial difference between oLT and iMT libraries (Additional file 1: Table S2). In addition, the expected nucleosome banding patterns were observed in the fragment size distribution for both oLT and iMT libraries (Fig. 1b). When we applied nucleoATAC to infer genome-wide nucleosome occupancy and positioning from ATAC-seq data [31], we found similar aggregated signals around TSS for both oLT and iMT libraries, corresponding to –2, – 1, +1, +2, +3 nucleosomes as well as nucleosome depletion region at upstream of TSS (Fig. 1c). Thus, we concluded that our ATAC-seq libraries had good and indistinguishable quality between oLT and iMT regions.

### Accessible chromatin landscape highlights potential enhancers and their target genes relevant to OA

Based on the 16 ATAC-seq libraries, we identified a set of unified accessible chromatin regions across all samples (n=109,215 robust peaks, Methods, Additional file 1: Table S3); 77,655 (71.1%) of which were annotated as enhancers and 18,410 (16.9%) as promoters (Fig. 2a) based on Roadmap DHS annotations (Methods). To assess the relevance of these peaks to OA, we intersected these peaks against Roadmap DHS, OA GWAS SNPs and OA DML datasets. We found that both OA GWAS SNPs and OA DML are significantly enriched in these peaks compared to all Roadmap DHS (GWAS SNP: Odds ratio = 4.3, p < 2 × 10^−16^; DML: Odds ratio = 1.8, p < 2 × 10^−16^, Fisher’s exact test), suggesting the peaks identified in this study are relevant to OA. In fact,majority of the OA GWAS SNPs (75.49%) that overlap our ATAC-seq peaks fall within annotated enhancers, which is consistent with the fact that GWAS SNPs reside in enhancers [14,52]. In this way, we identified 140 enhancers overlapping the OA GWAS SNPs (Additional file 1: Table S4) and 727 enhancers overlapping the OA DML (Additional file 1: Table S5). To further characterize these enhancers potentially relevant to OA, we utilized public resources [25,26] to predict their target genes (Methods). These enhancers and their predicted target genes represent candidates of dysregulated transcriptional network during OA pathogenesis. This included the previously identified OA associated genes, such as *FGFR3* [53,54] (its enhancer overlaps an OA GWAS SNP, Fig. 2b) and *PTEN* [55] (its predicted enhancer overlapsN with an OA DML, Fig. 2c), thus validating our approach. In addition, rs4775006 is a proxy SNP of rs4646626 (r^2^ = 0.816), which has previously been identified as a suggestive OA risk locus (p = 9 × 10^−6^, [56]). We found this SNP hitting the accessible enhancer region (chr15:57922910-57924140) in our samples, and predicted to target the *ALDH1A2* gene. Therefore, this study provides an accessible chromatin landscape of cartilage tissue, which allows better interpretation of other genetic and epigenomic data relevant to OA and other skeletomuscular disease related to cartilage tissue.

### Identification of differentially accessible enhancers in OA

To compare the accessible chromatin landscape between the intact (oLT) and the damaged (iMT) tissues, we performed principal component analysis on the peak signals across the 16 samples. The first principal component (41.37% of variance) can be attributed to tissue damage variations (i.e. oLT versus iMT), while the second principal component (16.03% of variance) can be attributed to patient-to-patient variations (Fig.3a). This observation suggests the accessible chromatin landscapes of damaged and intact tissues are readily distinguishable from each other, despite the variations among individual patients.

To identify the chromatin signatures relevant to OA, we performed differential accessibility analysis between oLT and iMT using edgeR [37]. First and foremost, we assessed the global relevance of the differential accessibility of peaks to OA. In Fig. 3b, we investigated the relationship between the extent of differential accessibility of peaks (measured by FDR) and their 1) compositions, 2) enrichment in OA GWAS SNPs, and 3) enrichment in OA DML. We found that the peaks that are more differentially accessible (i.e. smaller FDR) contain substantially higher fraction of enhancers (Fig. 3b, *top panel*), suggesting enhancers are more likely to be dysregulated than promoters during OA disease progression. Moreover, peaks that are more differentially accessible tend to be more enriched (i.e. higher odds ratio) in both OA GWAS SNPs and DML

(Methods, Fig. 3b, *middle and lower panels*). The differentially accessible enhancers overlapping either OA GWAS SNP or OA DML are summarized in Table 1 and 2, respectively. As a negative control, we also examined the GWAS SNPs and DML associated with Parkinson’s disease [43], which is a neurodegenerative disease pathologically unrelated to cartilage, and did not observe any substantial enrichment. Taken together, these observations suggest the differential chromatin accessibility measured in this study is relevant to OA disease progression in a specific manner.

**Table 1.** Differentially accessible enhancers overlapping with OA GWAS SNPs

| Enhancer peak ID | Proxy SNP ID | Lead SNP ID | r<sup>2</sup> | GWAS literature | Log<sub>2</sub> (fold change) | FDR | Predicted target gene(s) |
| --- | --- | --- | --- | --- | --- | --- | --- |
| chr15:57922910-57924140 | rs4775006 | rs4646626 | 0.816 | [56] | -0.546 | 0.050 | ALDH1A2 |
| chr3:46705071-46706482 | rs7426919 | rs6442021 | 0.835 | [40] | 0.581 | 0.023 | ALS2CL; PRSS45 |
| chr7:32616344-32617105 | rs10951345 | rs7805536 | 0.901 | [56] | -0.769 | 0.028 | AVL9 |
| chr7:32611048-32612186 | rs10807862 | rs7805536 | 0.901 | [56] | -0.957 | 0.006 | AVL9; LSM5 |
| chr8:8680144-8681063 | rs7837620 | rs1876836 | 0.908 | [56] | -0.804 | 0.037 | CLDN23; ERI1; MFHAS1 |
| chr8:8680144-8681063 | rs7837620 | rs1876836 | 0.908 | [56] | -0.804 | 0.037 | SGK223 |
| chr3:15275408-15276316 | rs4685243 | rs3773475 | 0.581 | [83] | 1.060 | 0.009 | SH3BP5 |
| chr15:34841146-34841782 | rs17237191 | rs717941 | 1 | [84] | -0.696 | 0.041 | ZNF770 |

**Table 2.** Differentially accessible enhancers overlapping with OA Differentially Methylated Loci

| Enhancer peak ID | DML | Delta beta | Log<sub>2</sub> (fold change) | FDR | Predicted target gene(s) |
| --- | --- | --- | --- | --- | --- |
| chr1:19398124-19398841 | cg06360604 | -0.177 | 1.194 | 0.003 | AKR7A2; AKR7A3; CAPZB; PQLC2 |
| chr16:66511371-66512170 | cg02934719 | -0.168 | 0.827 | 0.021 | BEAN1; TK2 |
| chr11:74025533-74026295 | cg01117339 | -0.164 | 0.623 | 0.037 | COA4; RAB6A; UCP2 |
| chr1:85606965-85607684 | cg00418071 | -0.157 | 1.214 | 0.001 | CYR61; DDAH1; ZNHIT6; SYDE2 |
| chr4:158809913-158811140 | cg11637968 | -0.218 | 0.878 | 0.032 | ETFDH; FAM198B; FNIP2; PPID; C4orf46 |
| chr5:32767956-32768626 | cg26647771 | -0.198 | 0.924 | 0.010 | GOLPH3; NPR3; TARS |
| chr1:226708432-226710048 | cg04503570 | -0.166 | 0.500 | 0.046 | PSEN2 |
| chr2:1720779-1722443 | cg08216099 | -0.193 | 0.732 | 0.016 | PXDN |
| chr10:119523073-119524950 | cg18591136 | -0.152 | 0.508 | 0.040 | RGS10 |
| chr6:168412123-168412714 | cg26379705 | -0.174 | 0.736 | 0.009 | SMOC2 |
| chr6:146850220-146851275 | cg18291422 | -0.184 | 0.960 | 0.012 | STXBP5 |
| chr8:81688679-81689538 | cg04491064 | 0.164 | -0.959 | 0.046 | CHMP4C |
| chr12:124700740-124701544 | cg04658679 | 0.160 | -0.514 | 0.030 | FAM101A; UBC |
| chr4:26412447-26413348 | cg22992279 | 0.159 | -0.737 | 0.028 | RBPJ |
| chr11:35344856-35345647 | cg13971030 | 0.157 | -0.656 | 0.038 | SLC1A2 |
| chr12:132846887-132848225 | cg21232015 | 0.151 | -0.615 | 0.038 | ZNF10; ZNF268; ZNF605; ZNF84; GOLGA3; ANKLE2; CHFR |

We next determined the significantly differentially accessible peaks with FDR ≤ 0.05. Out of the 4,450 differentially accessible peaks, 1,565 are more accessible and 2,885 are less accessible in the damaged tissues compared to the intact tissues (Fig. 3c, Fig. 3d, and Additional file 2: Figure S2a). The identified differentially accessible peaks cluster in two groups (Additional file 2: Figure S2b), and are generally consistent among patients (Additional file 2: Figure S2c). Majority (85.3%) of these differentially accessible peaks are enhancers and only 7.6 % are promoters. It is noted that the promoter accessibility of *SOX11* and *RGR* are altered in the OA damaged tissues (more accessible for *SOX11* and less accessible for *RGR*, Fig. 3d), which are consistent with the previous findings that they are up- and down-regulated in OA damaged tissues, respectively [20].

Majority of the differentially accessible peaks are enhancers, which are known to regulate transcription. Cell type-specific enhancers largely drive the transcriptional program required to carry out specific functions for each cell type [57] and their dysregulation may lead to diseases [58]. To assess the cell type specificity of the differentially accessible enhancers identified in this study, we examined their enrichment of cell type-specific enhancers of 125 cell types defined by the Roadmap Epigenomics Project [24]. We found these differential accessible enhancers are highly enriched for enhancers that are specific to bone-related cell types in damaged tissues (e.g. chondrocyte and osteoblast), as well as mesenchymal stem cells (Fig. 3e). This observation is consistent with the hypothesis that inappropriate activation of endochondral ossification process might play a role in OA progression [59], which is thought to occur through differentiation of mesenchymal stem cells to chondrocytes or osteoblasts, as well as hypertrophic differentiation of chondrocytes [60].

In summary, we demonstrate that the differential accessible chromatin regions identified in this study are enriched in OA-relevant enhancers supported by multiple genetic and epigenome evidence. These differentially accessible enhancers and their predicted target genes could be used for prioritization of candidate genes to be tested for studying OA disease progression.

### Motif enrichment analysis reveals transcription factors relevant to OA

In order to gain more insights into which regulatory pathways may be dysregulated in OA, we next examined the enrichment of transcription factor binding motifs in the differentially accessible regions. Enrichment analyses were performed separately for the regions that are significantly more or less accessible in the damaged tissues, using all accessible regions as background. Transcription factors with significantly enriched motifs are summarized (Additional file 2: Figure S3b), and their binding prediction in robust peaks are listed (Additional file 1: Table S3). We note most of these transcription factors belong to ETS and bZIP family; many of which are known to regulate genes involved in bone or cartilage development, including AP-1 [61], CEBP [62], MafK [63], STAT3 [64], and ERG [65,66]. Moreover, we have previously proposed that ETS-1 may be involved in OA based on our DML study [19]. Taken together, the motif Menrichment analysis of the differentially accessible regions in the damaged tissues is consistent with the hypothesis that the transcriptional program for chondrocyte differentiation may be disrupted during OA progression, and suggests that cell type-specific enhancers may be dysregulated through the ETS and bZIP family transcription factors.

### Integrative transcriptomics and epigenomics analysis reveals pathways involved in OA

To evaluate the effects of the dysregulated regulatory regions on gene expression in OA, we reanalyzed the RNA-seq dataset from a different cohort that used the same disease model (i.e. oLT vs. iMT) [15] and integrated it with the differentially accessible regions identified in this study (Fig. 4a). We first verified that the dysregulated regulatory regions have detectable effects on the gene expression; the genes with more accessible promoters or enhancers have significantly higher expression fold-changes between oLT and iMT (p < 0.001, Student’s *t-*test) than the ones with less accessible promoters or enhancers (Fig. 4b). To further investigate the congruence between these transcriptomic and epigenomic changes, we overlapped the significant differentially expressed genes (n=3,293, FDR ≤ 0.05) onto the genes with differentially accessible promoters (n=255) or enhancers (n=2,406) (Fig. 4c). We find that their overlaps are statistically significant (p < 0.001 in both promoters and enhancers, Fisher’s exact test), which further demonstrate the chromatin accessibility dataset from this study is generally consistent with the transcriptomic dataset. As a result, we identified 371 genes that are consistently dysregulated both at the epigenomic and transcriptomic levels, representing a shortlist of OA-related genes candidates supported by multiple lines of evidence (Fig. 4c, Additional file 1: Table S6).

**Fig. 4.**
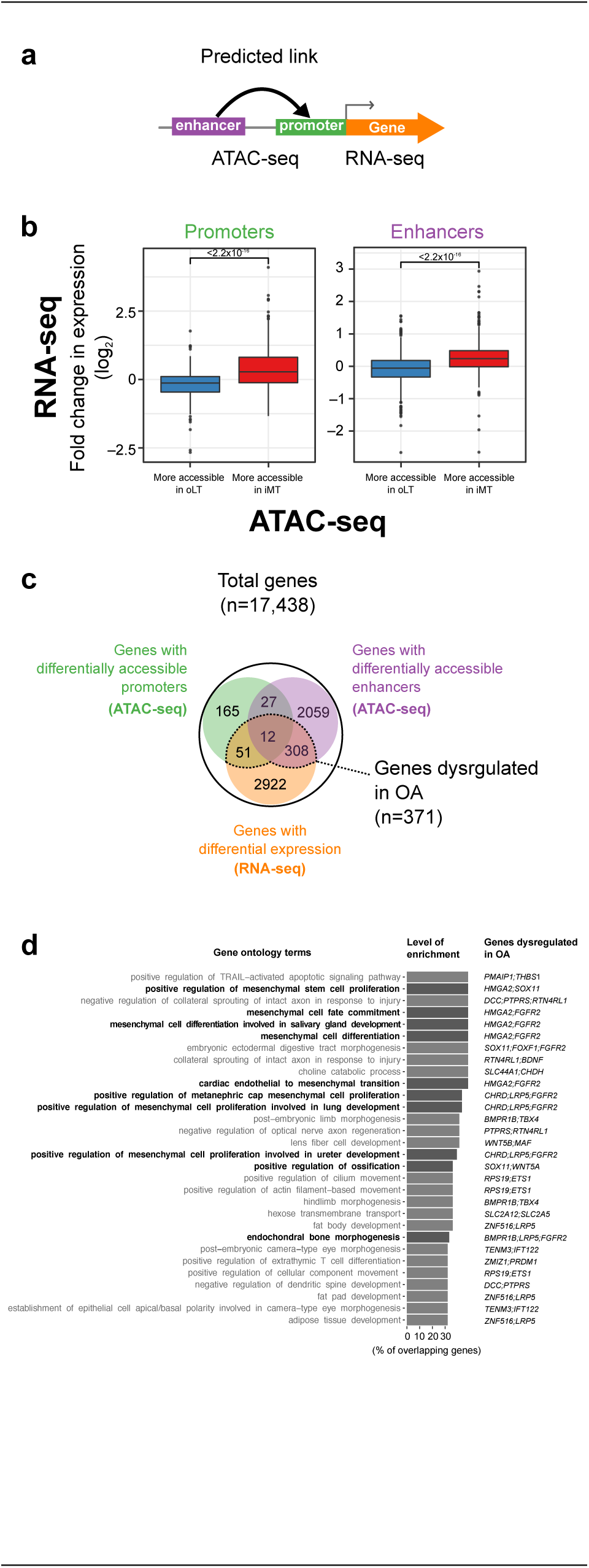
Integrative transcriptomic and epigenomic analysis reveals dysregulated genes and pathways in OA. (a) A scheme of ATAC-seq and RNA-seq integration analysis. (b) Differential chromatin accessibility (ATAC-seq) at both promoter (*left*) and enhancer (*right*) in OA is generally consistent with differential expression (RNA-seq [15]). Fold change between iMT and oLT is plotted. Box plots show the median, quartiles, and Tukey whiskers. (c) Venn diagram summarizing protein coding genes that are dysregulated both at the transcriptomic (RNA-seq) and epigenomic (ATAC-seq) levels in OA. (d) GO enrichment analysis of 371 genes from (c). Top 30 terms in GO biological process ranked by the level of enrichment and overlapping dysregulated genes are listed. Terms related to MSC, bones, and chondrocytes are highlight in bold.

To elucidate the biological pathways dysregulated in OA, we examined the enrichment of gene ontology (GO) terms of the 371 OA-related candidate genes using enrichR [48] (Figure 4d, top 30 terms listed). Overall, we observe the enrichment of GO terms related to cell fate and differentiation, including MSC differentiation, ossification and bone development (Fig. 4d, dysregulated genes in each GO terms are listed in Additional file 1: Table S7). In the ‘positive regulation of ossification pathway’, two genes were identified to be more accessible and upregulated: *SOX11* (only the promoter is more accessible) and *WNT5A* (both the promoter and the enhancer are more accessible) (Additional file 2: Figure S3b). It has been shown that WNT5A protein can induce matrix metalloproteinase production and cartilage destruction [67], and its upregulation is consistent with ossification being an important process and signature of OA progression. Other susceptible genes and pathways that support the ossification during OA from this analysis include *LRP5, FGFR2* and *BMPR1B* in the endochondral bone morphogenesis pathway, which have been reported as OA associated genes previously [68–70]. In conclusion, our integrative analysis of ATAC-seq and publicly available RNA-seq datasets indicates that dysregulated chondrocyte differentiation and endochondral ossification are central to OA progression.

## Discussion

Conventional epigenomic profiling at the chromatin level, such as chromatin immunoprecipitation sequencing (ChIP-seq) and DNase-seq, is informative in providing insights into the molecular mechanism underlying the regulation of gene expression. However, applying these methods to clinically relevant tissue is less feasible due to the requirement of large number of cells. Especially with the cartilage tissues, the limited tissue sampling size and the extracellular matrix make collection of sufficient cells difficult. One major advantage of ATAC-seq is that it can be achieved with only thousands of cells, making the direct chromatin profiling of clinical samples feasible. In this study, we applied ATAC-seq on OA samples to obtain a chromatin accessibility map in articular cartilage, and identified regulatory regions associated with OA. The lack of normal knee tissues due to difficulties associated with collecting them is a limitation for studying OA. However, our previous studies of the pathology andM transcriptome showed that the oLT regions are very similar to normal [16,20].Thus, the oLT regions could serve as a suitable alternative to normal control, which could also reduce the inter-individual variations.

Our strategy of integrating the epigenomic data of clinically relevant tissues with the publicly available genetic and transcriptomic data allowed us to better understand how the identified loci may contribute to OA pathogenesis. Most of these accessible chromatin regions are annotated enhancers and we linked them to their putative target genes using public datasets. With this enhancer-gene map in chondrocyte, we can now better interpret the previously identified OA GWAS SNPs or OA differential methylated loci located lie outside of the coding regions. For example, we have verified that the previously reported OA associated SNP rs4775006 is indeed accessible in chondrocytes, that lies within the enhancer region of the predicted target gene *ALDH1A2.* Since *ALDH1A2* is inactivated in prechondrogenic mesenchyme during the cartilage development [71], this SNP may contribute to OA through disrupting the enhancer of *ALDH1A2* and inappropriately activating a cell differentiation pathway. In addition, we identified several aberrantly methylated enhancers that may be associated with OA. One example is cg09221159 within the enhancer for *PTEN* gene which is hypo-methylated in damaged cartilage [19]. It is involved in the positive regulation of the apoptotic signaling pathway and has been proposed that chondrocyte apoptosis contributes to the failure in appropriately maintaining the cartilage. Thus, our analyses show that the chromatin accessibility map can provide an additional layer of evidence for determining which loci, especially those in the non-coding regions, are associated with OA, which may have been ignored in previous studies.

In general, our differential enhancer analysis shows MSC, chondrocyte and osteoblast-specific enhancers are dysregulated in the damaged tissues. Furthermore, motif enrichment analysis of differentially accessible loci has identified many dysregulated transcription factors, the functions of which are known to be in chondrocyte development regulation. For example, the transcription factor Jun-AP1, which is enriched in the more accessible regions of the damaged tissues, is known to regulate chondrocyte hypertrophic morphology, which contributes to longitudinal bone growth, consistent with the notion of endochondral ossification [72]. A recent study showed injection of adipose-derived stromal cells overexpressing an AP-1 family transcription factor Fra-1 can inhibit OA progression in mice [73]. In our ATAC-seq analysis, other members of AP-1 family (e.g. BATF, FOSL2) also showed enrichment in differentially accessible regions, which could be candidate targets for such experiments.

In the integrative analysis of ATAC-seq and RNA-seq, many dysregulated genes related to lineage differentiation of MSC pathways were observed. For example, we found that *BMPR1B* (bone morphogenetic protein receptor type 1B) is upregulated and both its promoter and enhancer are more accessible in the damaged tissue. Its activation is consistent with the ossification pathway activation, since it encodes a transmembrane serine kinase that binds to BMP ligands that positively regulate endochondral ossification and abnormal chondrogenesis [74,75]. Consistently, an osteoblast marker gene *MSX2* (Msh Homeobox 2) involved in promoting osteoblast differentiation [76,77], is upregulated and both its promoter and enhancer are more accessible in the damaged tissue, suggesting the osteoblast differentiation may be activated in OA. Furthermore, we found *ROR2* (receptor tyrosine kinase like orphan receptor 2) is down-regulated and its enhancers are less accessible in the damaged samples. Since it is required for cartilage development [78,79], it suggests that the normal chondrocyte development and cartilage formation may be compromised in OA. Consistently, we have also determined *FGFR2* and *STAT1*, which are known to inhibit chondrocyte proliferation [80], are upregulated. In summary, our ATAC-seq data and integrative analyses support the OA model of abnormal MSC differentiation and endochondral ossification over other models, such as inflammatory mediums from the synovium.

The pathogenesis for OA is not yet fully understood, despite multiple genes and pathways that have been characterized to be dysregulated [13,15,42,81,82]. In this study, by integrating clinically relevant epigenomic data with genetic and transcriptomic data, we provide multiple lines of evidence supporting a number OA candidate genes and pathways that may be crucial to OA pathogenesis, which could potentially be used for clinical diagnostic or as therapeutic targets.

## Conclusions

We present in this work an application of ATAC-seq in OA in a clinical relevant setting. The chromatin accessibility map in cartilage will be a resource for future GWAS and DNA methylation studies in OA and other musculoskeletal diseases. We identified altered promoters and enhancers of genes that might be involved in the pathogenesis of OA. Our analyses suggest aberrant enhancer usage associated with MSC differentiation and chondrogenesis in OA. Understanding these molecular basis of OA is necessary for future therapeutic intervention.

## Abbreviations

ATAC-seq: Assay for Transposase-Accessible Chromatin with high throughput sequencing
CPM: count-per-million
DHS: DNase I Hypersensitive Sites
DML: Differentially Methylated Loci
eQTL: expression Quantitative Trait Loci
FDR: False Discovery Rate
GWAS: Genome-wide association studies
iMT: inner region of medial tibial plateau
MSC: Mesenchymal Stem Cell
OA: Osteoarthritis
oLT: outer region of lateral tibial plateau
SNP: Single Nucleotide Polymorphisms
TSS: Transcription Start Sites

## Ethics, consent and permissions

This study was approved by the ethics review board of all the participating institutions (Sagamihara National Hospital and RIKEN).

## Consent to publish

Informed consent was obtained from each patient enrolled in this study.

## Availability of data and materials

Raw and processed data generated are available on the Gene Expression Omnibus under the accession code GSE108301. [Reviewer access code: cpynocyaxnsvfsr]

## Competing interests

All authors declare that they have no conflict of interest.

## Funding

This work was supported by a Research Grant from the Japanese Ministry of Education, Culture, Sports, Science and Technology (MEXT) to the RIKEN Center for Life Science Technologies, RIKEN Epigenetics Program, and Ye Liu was supported by the RIKEN Junior Research Associate Program.

## Authors’ contributions

The study was conceived and designed by AM and MTML. Sample collection were done by NF and NT. Experiments were performed by YL. Analysis and interpretation of data were carried out by YL, JC and CH. The study was supervised by AM, MTMKL and ZZ. YL and JC drafted and AM, CH and MTML revised the manuscript. All authors read and approved the final version of the manuscript.

## Acknowledgements

We thank Dr. Shiro Ikegawa (RIKEN & Tokyo University) for his support on the project. We thank Yanfei Zhang (Geisinger), Tommy Terooatea (RIKEN), Mitsunori Yahata (RIKEN), Haruka Yabukami (RIKEN) and Prashanti Jeyamohan (RIKEN) for active discussions, technical assistance and support. We thank GeNAS for the sequencing service.

## Additional files

Additional file 1: Supplemental tables S1-S7. (XLSX, 12.8 MB)

Additional file 2: Supplemental figures S1-S3. (PDF, 870 KB)

## Figure legends

**Additional file 2: Figure S1. ATAC-seq of human cartilage.** (a) Images of the dissected cartilage tissue from fresh articular tibial bone for all patients. Section regions of tibial plateau: outer region of lateral tibial plateau (oLT) exhibited macroscopically normal cartilage with a visibly smooth cartilage surface, and inner region of medial tibial plateau (iMT) had visible severe erosion of cartilage. (b) Enrichment qPCR of a housekeeping gene (*GAPDH*) over a heterochromatin region is correlated with enrichment of ATAC-seq signal in DHS regions.

**Additional file 2: Figure S2. Differential ATAC-seq peaks. (**a) Mean-difference plot of all ATAC-seq peaks. Red and blue dots represent peaks that are more- and less-accessible in iMT, respectively. (b) Principal Component Analysis of differential ATAC peaks between oLT (*blue*) and iMT (*red*). (c) Genome browser views showing consistency of ATAC-seq signals across patients at example loci.

**Additional file 2: Figure S3. Data integration.** (a) Top *de novo* motif (*top*) and top predicted known transcription factors (*bottom*) enriched in differentially accessible regions. (b) An example gene (*WNT5A*) that is dysregulated at all three criteria (promoter accessibility, enhancer accessibility and expression).

**Figure S1.**
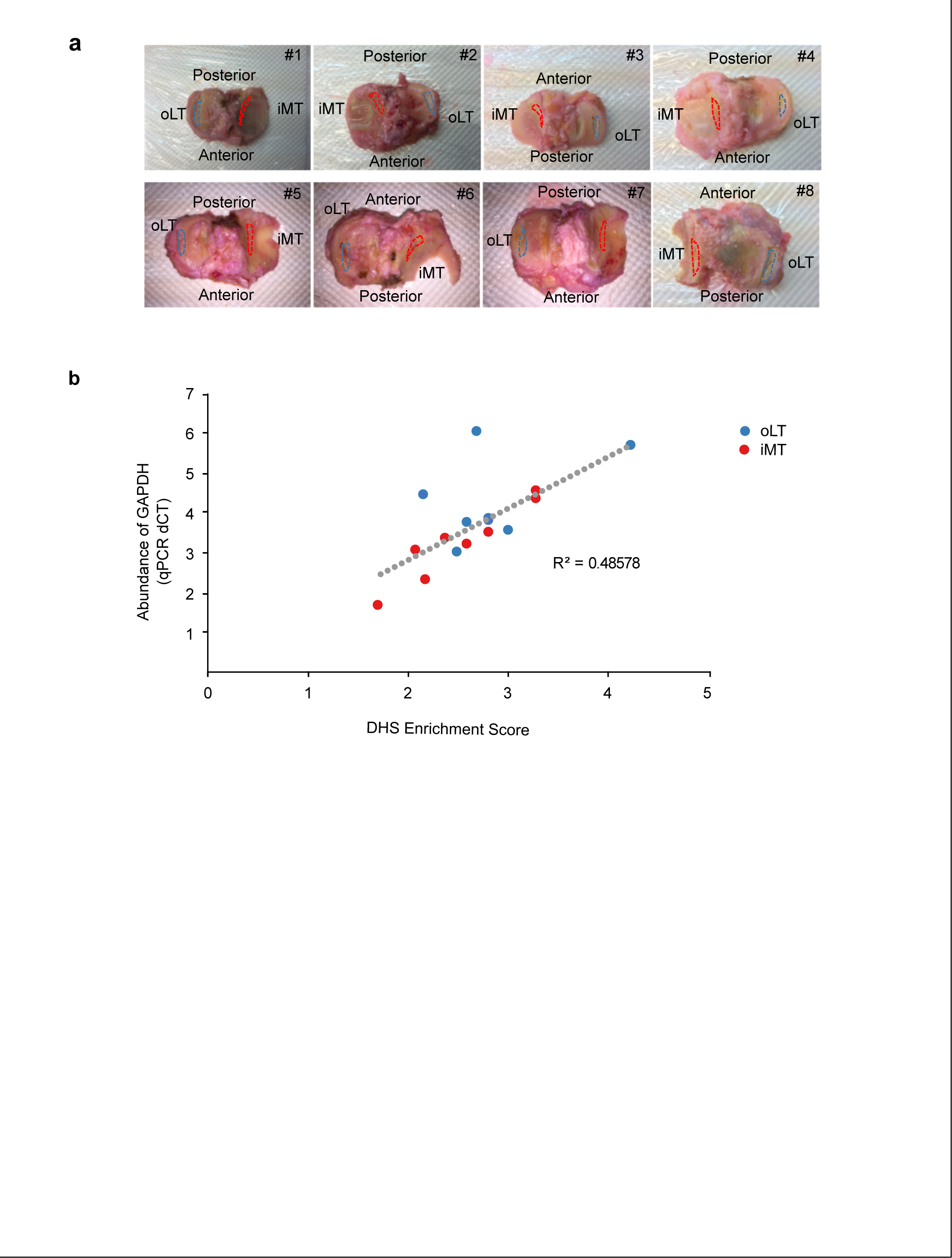

**Figure S2.**
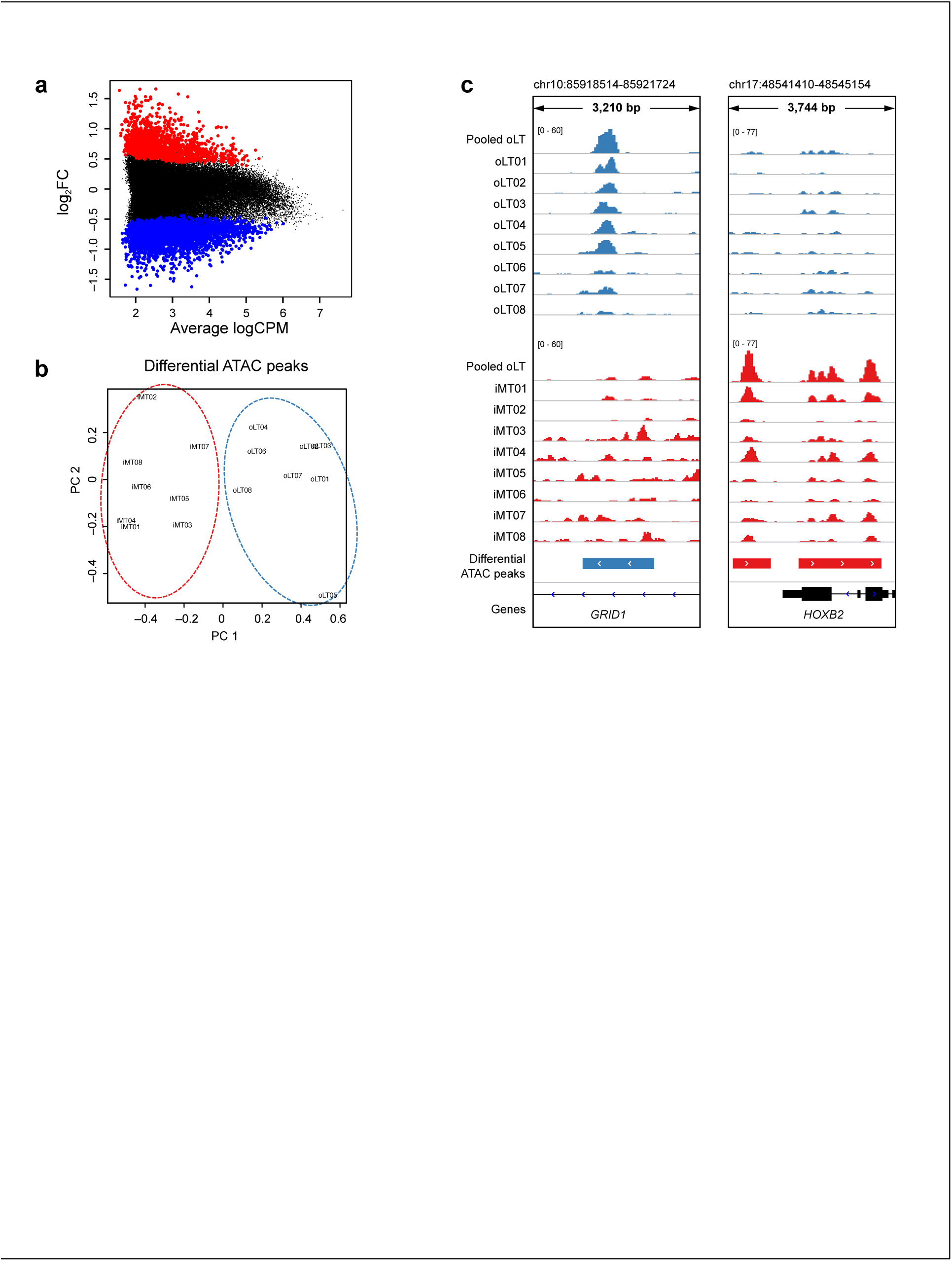

**Figure S3.**
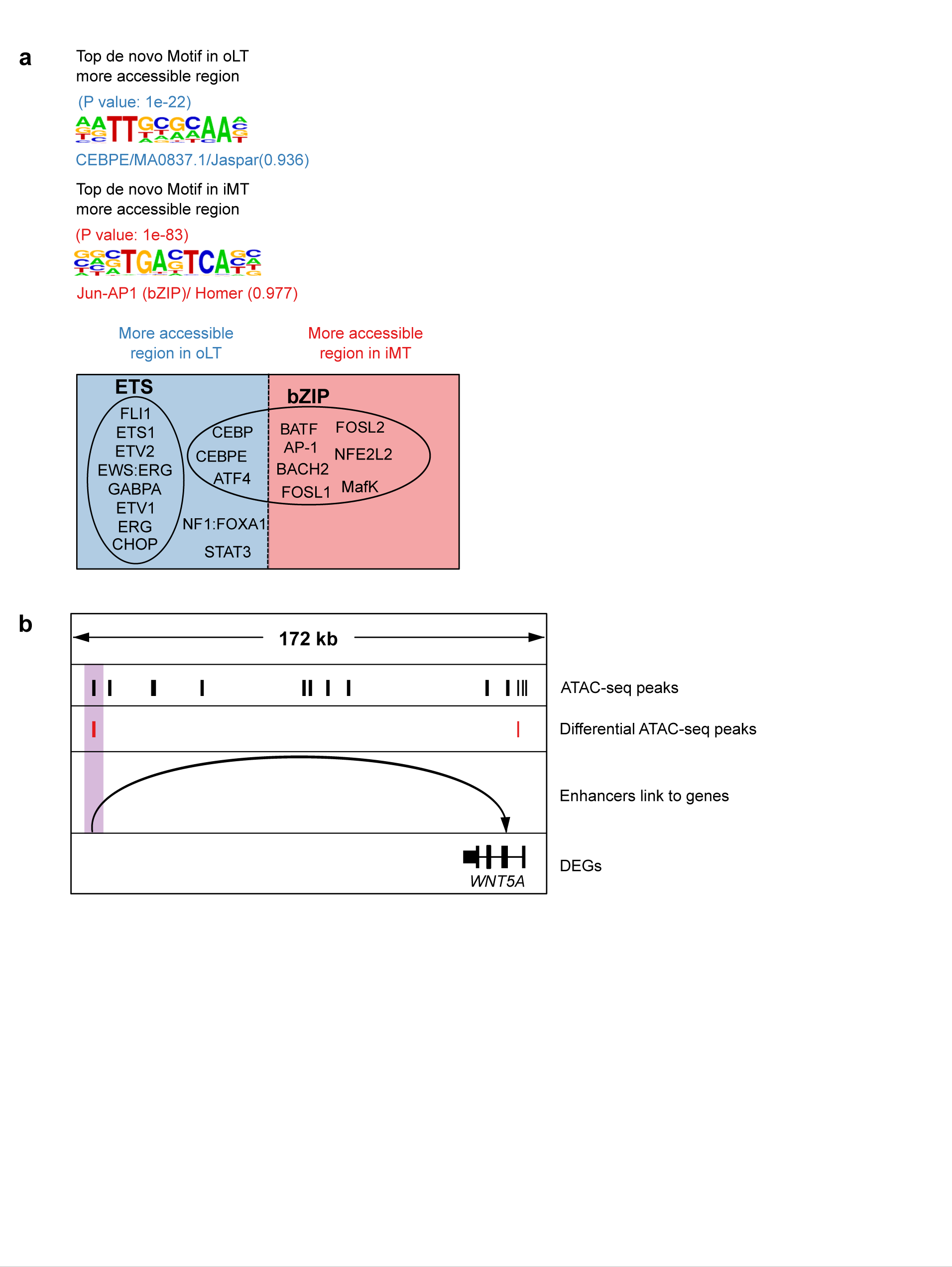

**Table 3.**
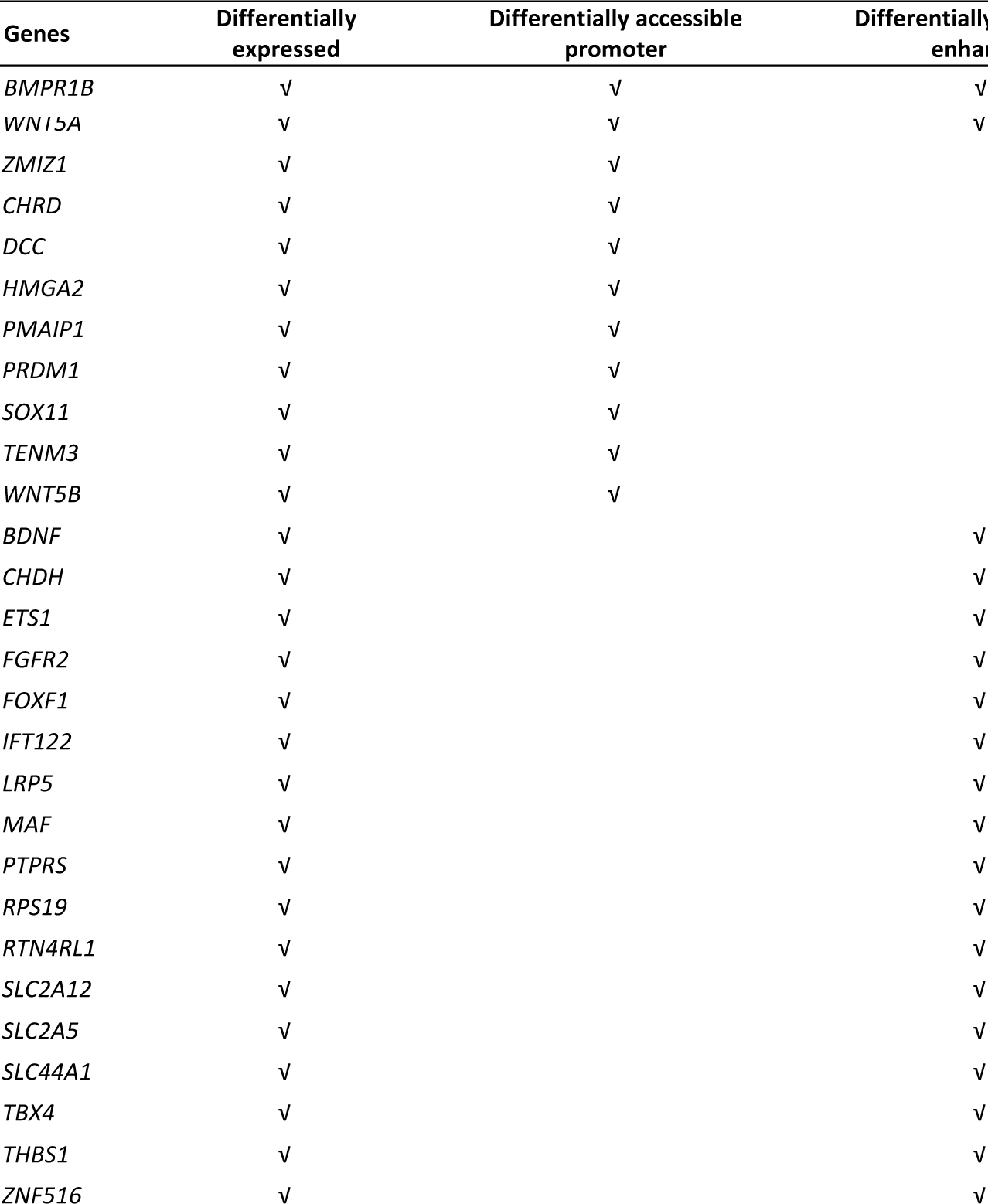
OA associated genes enriched in top 30 GO terms

